# Broca’s area responsible for speech production is regulated by lung functions

**DOI:** 10.1101/2024.11.20.624446

**Authors:** Siyu Cao, Wenwen Zhuang, Yuqian Hu, Shijun Qiu, Li-Hai Tan

## Abstract

For more than one and a half centuries, Broca’s area, specifically, the pars opercularis and pars triangularis in the left inferior frontal gyrus, has been known to be crucial for human speech production. However, it remains unanswered why this region is recruited for speaking. Speech production involves not only conceptualization and motor planning, but also respiration that provides the airflow necessary for creating sounds. Thus, the role of Broca’s area in speech may be shaped by the functionality and the related brain regions associated with lungs. To test this hypothesis, we recruited the patients suffering from chronic obstructive pulmonary disease (COPD) and required them to speak out words while scanning their brains with functional and quantitative magnetic resonance imaging (fMRI and qMRI). We observed that COPD patients exhibited altered cortical responses in left inferior prefrontal cortex and other regions during speech tasks, with abnormal activation in cortical sites associated with breathing. In addition, using the qMRI to generate longitudinal relaxation time (T_1_) maps as an index of brain microstructural changes including dendritic maturation and myelination, we discovered that the patients showed significantly higher T_1_ values than the control group in Broca’s area, suggesting reduced myelination and impaired microstructural integrity. Crucially, the data indicated that more severe dyspnea was associated with less well-developed microstructure in Broca’s area and its weaker activation. Our study has demonstrated for the first time that lungs may function to shape Broca’s area as the speaking center, and this is consistent with the recent work of a lung–brain axis.

## INTRODUCTION

Broca’s area, located in the left inferior frontal gyrus, is widely known as the speech production center discovered by Paul Broca in 1861. This brain area is adjacent to the precentral gyrus on the left, and assumed to be related to the motor planning and articulation stages to produce speech sounds^1-11^. Speech production, however, is a complex process that involves not only conceptualization and motor planning, but also respiration that provides the airflow necessary for speaking^12,13^. Air is drawn into the lungs and expelled through the trachea, a process regulated by the brainstem and bilateral inferior, middle frontal and premotor cortices^14,15^, which facilitate sound production. Therefore, Broca’s area and related brain regions may be shaped by the functionality of the lungs. This hypothesis is in harmony with the recent animal model findings of a lung–brain axis which indicate that the pulmonary microbiome is related to the central nervous tissue^16^.

To test this hypothesis with human participants, we recruited a group of patients suffering from chronic obstructive pulmonary disease (COPD) (Table S1) and asked them to overtly name commonly-used high-frequency words while scanning their brain activity and structure with magnetic resonance imaging (MRI). We contrasted their cortical activation and myelination alternation with normal healthy control participants, particularly focusing on Broca’s area and related cortical regions (such as the middle frontal gyrus and the precentral cortex) governing breathing. COPD is a progressive disease characterized by chronic irreversible airflow limitation in the lungs^17,18^, leading to the change of brain structure and function associated with breathing^19-21^. Thus, the COPD patients should exhibit altered cortical responses during naming performance, and cortical regions related to breathing function abnormally in speaking. This will constitute one of the mechanisms why lung function may be crucial to the speech function associated with Broca’s area.

## RESULTS

### Atypical Brain Activation in COPD patients

Since we asked our participants to speak out nouns and verbs in Chinese, which were compared with a baseline condition where the participants passively viewed non-word symbols, we first contrasted the brain activations of the two groups in each of the two experimental conditions (relative to the baseline). Activations were significantly weaker in COPD patients than in controls in naming nouns as well as verbs (Figure 1a and 1b, FDR-corrected, *P* < 0.05), and there was no reliable difference in brain activation between nouns and verbs (Figure S1), consistent with previous research^22,23^.

**FIGURE 1.**
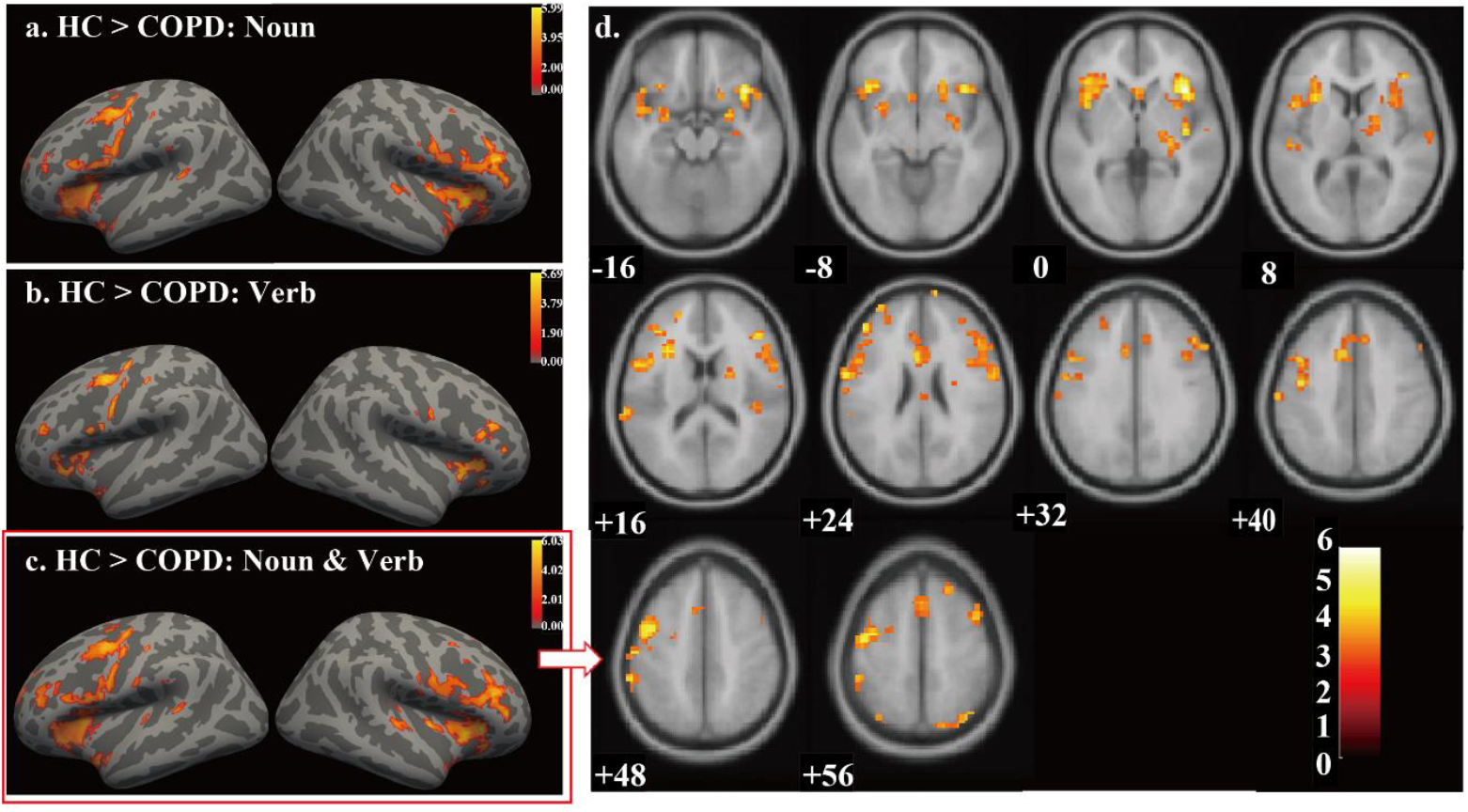
Statistical parametric maps of group activation differences between tasks (FDR-corrected *P* <0.05 with spatial extent n≥10 voxels). (a) represented the group differences during Noun contrasted with non-character symbols. (b) represented the group differences during Verb contrasted with non-character symbols. (c) and (d) represented the group differences during Noun and Verb conditions contrasted with non-character symbols.

Then we collapsed the data from nouns and verbs for further analyses. We focused on brain regions associated closely with naming. The activations in the COPD group were significantly weaker than that in the HC group (see Figure 1c, 1d, and Table S2) across multiple regions, including bilateral precentral cortex (BA 6; MNI: -45, 0, 45; 42, 6, 57), bilateral postcentral cortex (BA 43; MNI: -63, 0, 24; 60, -3, 24), and bilateral peri-insular cortex (BA 48; MNI: -42, 15, -3; 39, 18, 0). Importantly, we identified weaker activity in several key speech production areas in the COPD group, including bilateral inferior frontal gyrus (BA 44/45; MNI: -51, 9, 27; -45, 39, 21; 48, 27, 33; 45, 30, 18), bilateral middle frontal gyrus (BA 9; MNI: -45, 9, 45; 45, 15, 54) and bilateral precentral cortex (BA 6; MNI: -45, 0, 45; 42, 6, 57). The Cohen’s d values for these regions were all greater than 0.8, indicating a large effect size and suggesting a reliable difference in brain activation between the groups in these regions (see Table 1).

**TABLE 1.**
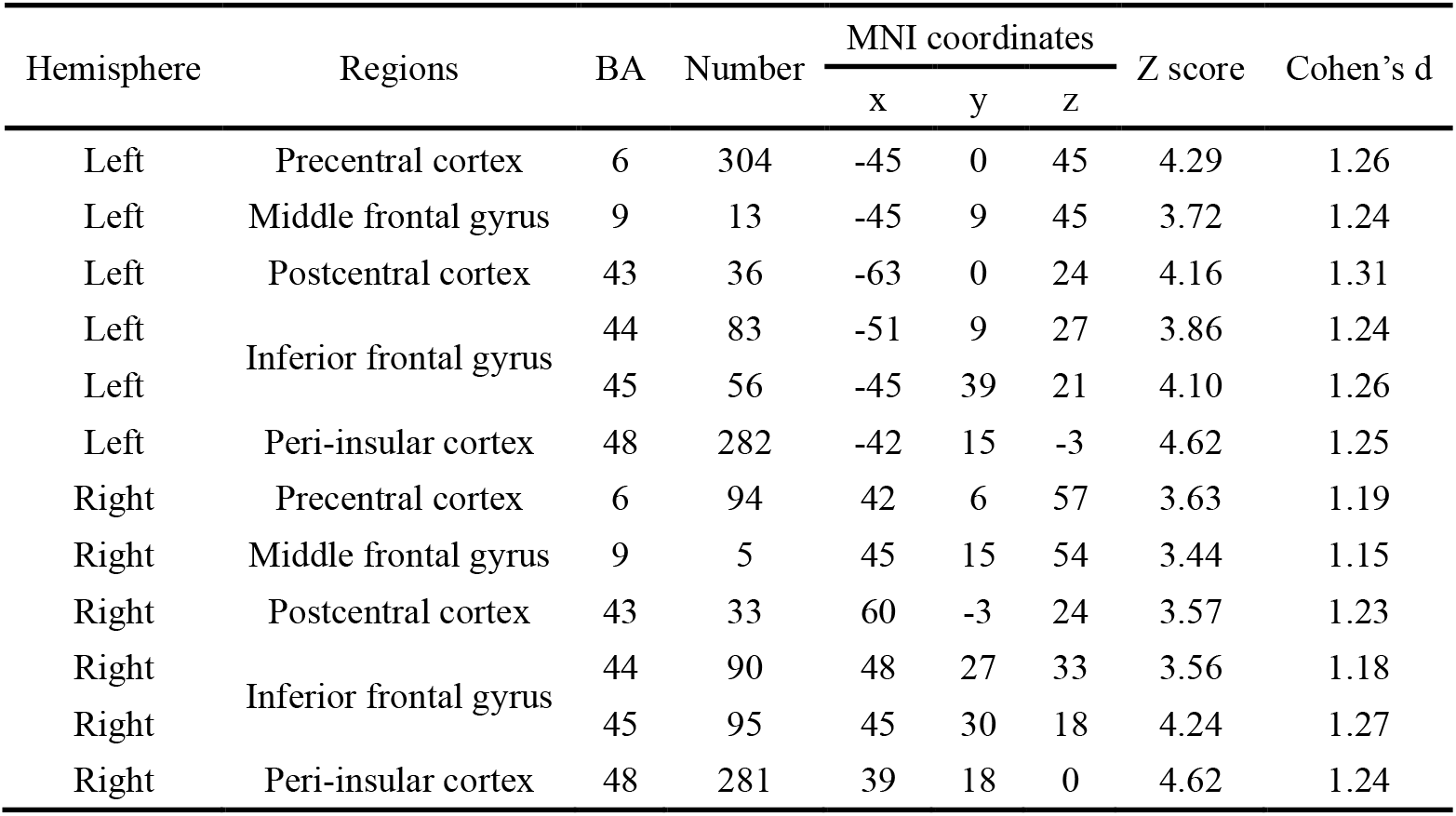
MNI Coordinates and activation statistics of the ROIs.

| Hemisphere | Regions | BA | Number | MNI coordinates |  |  | Z score | Cohen's *d* |
| --- | --- | --- | --- | --- | --- | --- | --- | --- |
|  |  |  |  | x | y | z |  |  |
| Left | Precentral cortex | 6 | 304 | -45 | 0 | 45 | 4.29 | 1.26 |
| Left | Middle frontal gyrus | 9 | 13 | -45 | 9 | 45 | 3.72 | 1.24 |
| Left | Postcentral cortex | 43 | 36 | -63 | 0 | 24 | 4.16 | 1.31 |
| Left | Inferior frontal gyrus | 44 | 83 | -51 | 9 | 27 | 3.86 | 1.24 |
| Left |  | 45 | 56 | -45 | 39 | 21 | 4.10 | 1.26 |
| Left | Peri-insular cortex | 48 | 282 | -42 | 15 | -3 | 4.62 | 1.25 |
| Right | Precentral cortex | 6 | 94 | 42 | 6 | 57 | 3.63 | 1.19 |
| Right | Middle frontal gyrus | 9 | 5 | 45 | 15 | 54 | 3.44 | 1.15 |
| Right | Postcentral cortex | 43 | 33 | 60 | -3 | 24 | 3.57 | 1.23 |
| Right | Inferior frontal gyrus | 44 | 90 | 48 | 27 | 33 | 3.56 | 1.18 |
| Right |  | 45 | 95 | 45 | 30 | 18 | 4.24 | 1.27 |
| Right | Peri-insular cortex | 48 | 281 | 39 | 18 | 0 | 4.62 | 1.24 |

Furthermore, we computed the correlation between the modified Medical Research Council (mMRC) Dyspnea Questionnaire scores and the activation values at the peak voxel coordinate in these ROIs. Our analysis revealed negative correlations between activation values in these regions and mMRC scores (see Table 2), particularly in the left inferior frontal gyrus (BA 44: *r* = -0.61, *P* = 0.00023; BA 45: *r* = -0.37, *P* = 0.037) and the left peri-insular cortex (*r* = -0.60, *P* = 0.00033).

**TABLE 2.** Correlations between the modified Medical Research Council (mMRC) Dyspnea Questionnaire scores and activation values at the peak coordinates in ROIs, as well as mean T1 values in left BA 44.

| Regions |  | Peak MNI coordinates |  |  | mMRC |
| --- | --- | --- | --- | --- | --- |
|  |  | x | y | z | *r* value |
| Activation values | Precentral cortex | -45 | 0 | 45 | -0.50** |
|  |  | 42 | 6 | 57 | -0.41* |
|  | Middle frontal gyrus | -45 | 9 | 45 | -0.41* |
|  |  | 45 | 15 | 54 | -0.47** |
|  | Postcentral cortex | -63 | 0 | 24 | -0.49** |
|  |  | 60 | -3 | 24 | -0.45** |
|  | Inferior frontal gyrus (BA 44) | -51 | 9 | 27 | -0.61*** |
|  |  | 48 | 27 | 33 | -0.38* |
|  | Inferior frontal gyrus (BA 45) | -45 | 39 | 21 | -0.37* |
|  |  | 45 | 30 | 18 | -0.35* |
|  | Peri-insular cortex | -42 | 15 | -3 | -0.60*** |
|  |  | 39 | 18 | 0 | -0.47** |
| mean T<sub>1</sub> value | Inferior frontal gyrus (BA 44) | - | - | - | 0.47** |

Note: \* $P < 0.05$ , \*\* $P < 0.01$ , \*\*\* $P < 0.001$ .

Speech production as a complex process is mediated by Broca’s area and regions responsible for articulatory gestures and auditory perception, which work together to coordinate the act of speaking and listening to ensure accurate speech output^9,24-27^. Consistent with this theory, we found that the COPD patients exhibited weaker activity in superior temporal gyrus (MNI: -66, -33, 15; 63, -18, 6) relative to the controls.

The COPD patients may have multiple neurocognitive problems that are not specific to speech production. To address this question, we analyzed the brain activities provoked by the baseline condition where the two groups of participants passively viewed non-words. We found no significant differences (FDR-corrected, *P* < 0.05; uncorrected, *P* < 0.001). In addition, the Montreal Cognitive Assessment indicated no reliable difference between the two groups (COPD: 19.56 ± 4.23; HC: 19.25 ± 3.89; *P* = 0.83, *t* = 0.22, see Table S1).

Therefore, we believe that our findings were specifically associated with speaking.

### ROIs for breathing based on the literature

Because the middle frontal gyrus, pre-supplementary motor area, inferior frontal gyrus, precentral gyrus and peri-insular area have been repeatedly demonstrated to be associated with breathing, we selected ROIs regions consistent with those reported in the published studies related to the lung function and conducted further analyses to examine whether there are regional differences in brain activation between HC and COPD participants. These ROIs included the bilateral pars opecularis (MNI: -54, 8, 16; 50, 10, 18)^28^, the left triangularis part of the inferior frontal gyrus (MNI: -50, 36, 22)^29^, the left inferior frontal gyrus (MNI: -30, 50, 20)^30^, the left middle frontal gyrus (MNI: -40, 40, 16)^29^, the right pre-supplementary motor area (MNI: 4, 12, 56)^28^, the left inferior precentral gyrus (MNI: -52, -8, 50)^31^, the left middle insula (MNI: -42, 2, 2)^30^ and the right insula (MNI: 38, 10, 8; 45, 16, -2)^32^.

We extracted the activation values at specified MNI coordinates for both groups and conducted a two-sample t-test. The results showed significantly higher activation in the HC group compared to the COPD group (see Figure 2). For instance, the left pars opecularis (MNI: -54, 8, 16), the left inferior frontal triangularis (MNI: -50, 36, 22), and the left inferior frontal gyrus (MNI: -30, 50, 20) all showed a marked increase in activation in the HC group compared to the COPD group. The data indicated that the COPD patients’ lung function was much weaker related to HC participants, which further led to hypoactivation in naming.

**FIGURE 2.**
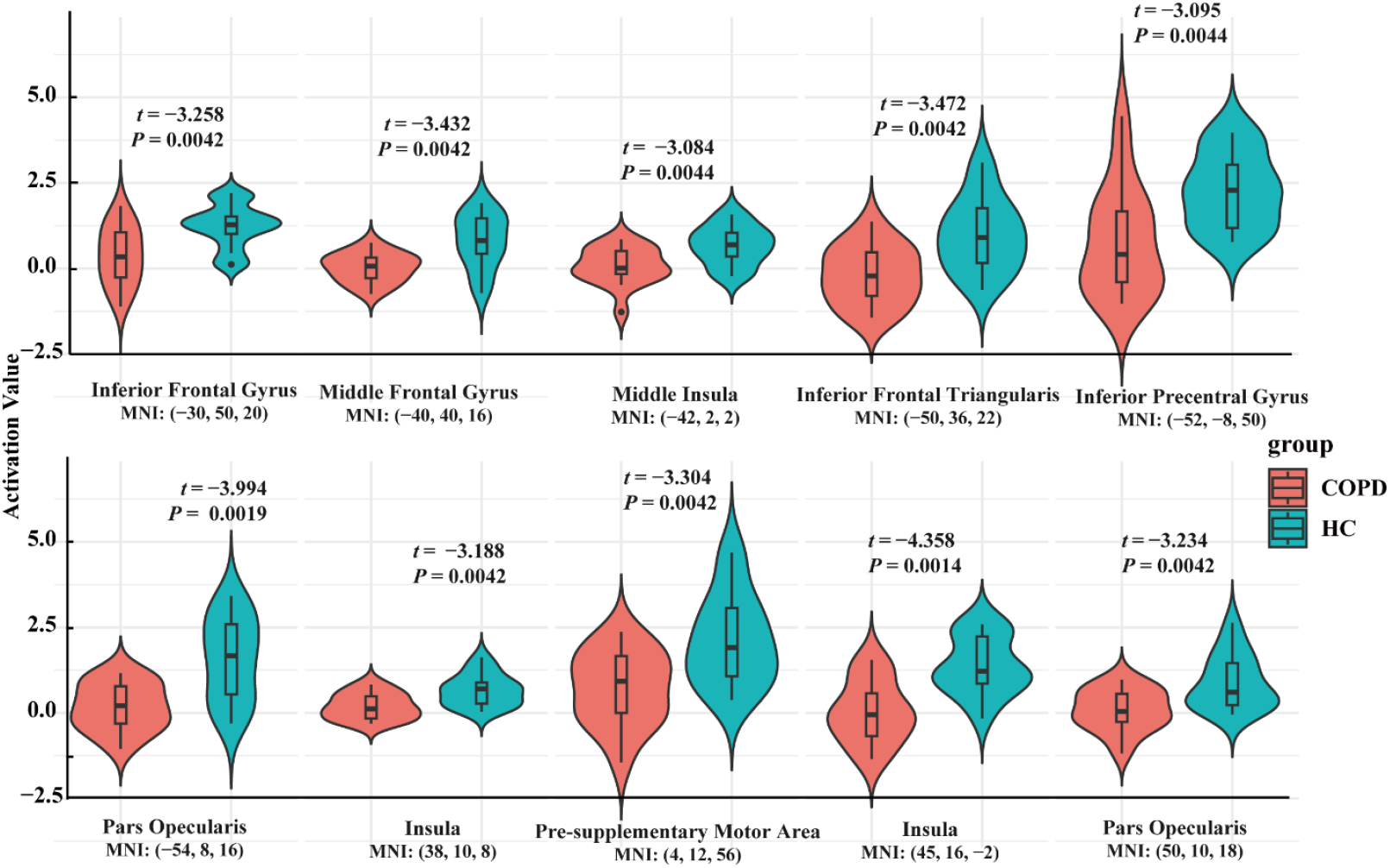
Activation statistics of literature-defined regions. Each plot corresponds to peak activation coordinates in MNI space, based on previously reported studies. Activation values were extracted from individual-level analyses comparing noun and verb conditions against baseline. The group differences were tested using two-sample t-tests. *P*-values were corrected for multiple comparisons using the Benjamini-Hochberg false discovery rate (FDR) method.

### Microstructure differences in T_1_

To quantify the cortical myelin architecture from a microstructural perspective, we employed quantitative MRI (qMRI) to generate longitudinal relaxation time (T_1_) maps, which are linearly related to iron and myelin concentrations^33-35^. The developmental decline in T_1_ is believed to reflect microstructural changes, such as dendritic maturation, myelination, and the proliferation of oligodendrocytes^36^. We calculated average T_1_ values within Broca’s area (left BA44) to assess group differences in tissue microstructure. Our findings indicated that the COPD group had significantly higher T_1_ values than the HC group (Figure 3, *P* = 0.0145, *Z* = 2.18). Cohen’s d for this region was 0.72, indicating a moderate effect size and suggesting a substantial difference between the groups in Broca’s area. As higher T_1_ values are typically associated with lower tissue density and reduced myelination, these results suggest reduced myelination and microstructural integrity in the COPD group.

**FIGURE 3.**
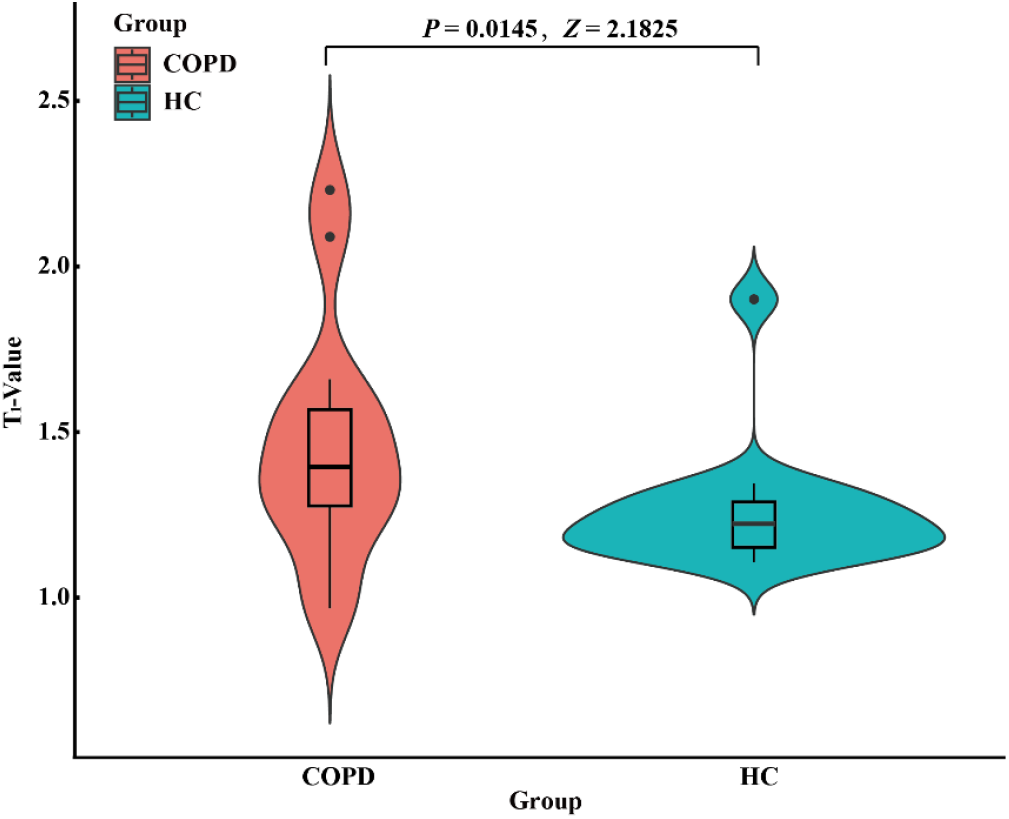
Statistical map of mean T_1_ values in left BA 44 comparing COPD and HC groups using a right-tailed Wilcoxon rank-sum test.

We computed the Pearson correlation coefficient between the mean T_1_ value in the left BA 44 and mMRC Dyspnea Questionnaire scores, finding a significant positive correlation (Table 2, *r* = 0.47, *P* = 0.0089). This suggested that higher T_1_ values were associated with greater mMRC scores, reflecting more severe dyspnea. After controlling for education level, we also found a negative partial correlation between mean T_1_ values and mean values of the top 30% activation values in the left BA 44 (Figure 4, *r* = -0.39, *P* = 0.0489). By contrast, in the baseline condition, no significant partial correlation was observed between the top 30% activation values and mean T_1_ values (*r* = -0.0052, *P* = 0.9799), suggesting that the observed negative correlation in the experimental condition is specific to task-related activations.

**FIGURE 4.**
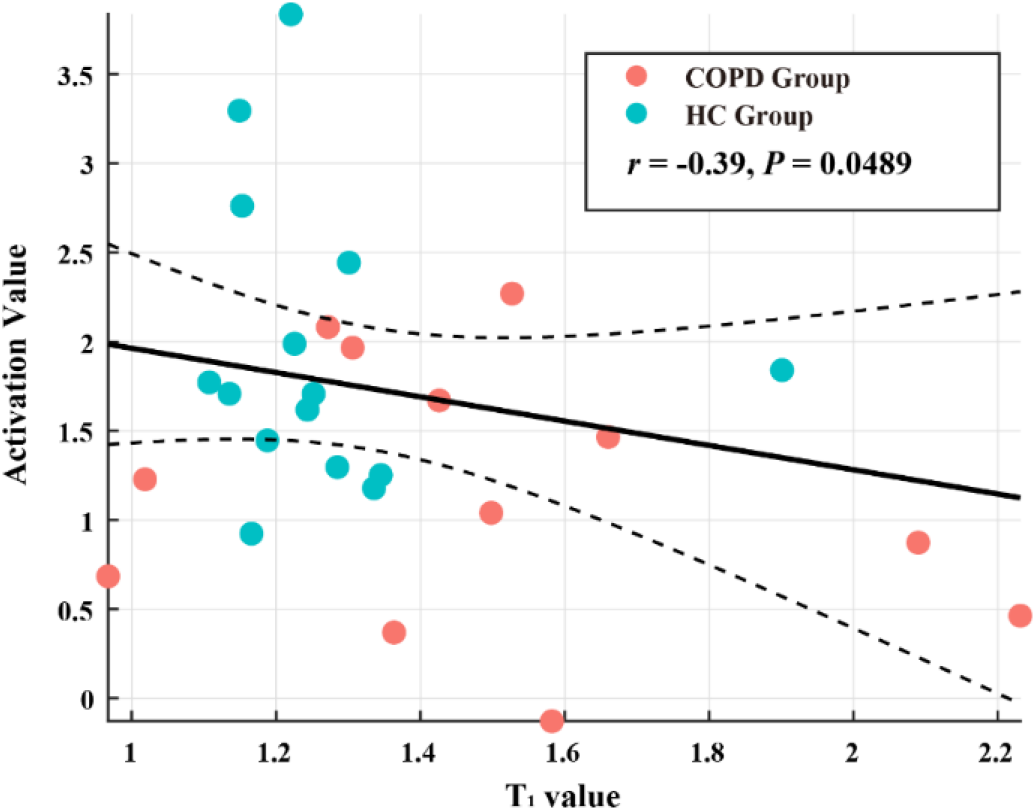
Partial Correlation between mean T_1_ values and mean values of the top 30% activation values in the left inferior frontal gyrus (BA 44), controlling for the influence of education level.

## DISCUSSION

Broca’s area located in the left inferior frontal gyrus is one of the most well-recognized brain regions supporting speech production. It is spatially close to the premotor cortex mediating motor coordination which is important for speaking. Since speaking engages respiration^8^, we assumed that cortical sites governing functions of lungs may functionally as well as structurally regulate Broca’s area. Our findings have provided important evidence for this hypothesis. Using a simple speaking task, we found that the brain areas such as the left inferior frontal gyrus (BA 44/45; MNI: -51, 9, 27; -45, 39, 21) and the precentral cortex (BA 6; MNI: -45, 0, 45) essential for speech production showed much weaker activities in the COPD patients. Crucially, we extracted literature-based ROIs related to lung functions and found that there were significant differences in activation between HC and COPD participants in the left pars opecularis (MNI: -54, 8, 16) and the left pars triangularis (MNI: -50, 36, 22), in addition to other regions, which control the functionality of lungs. These finding evidence that lungs function shapes Broca’s area as the speech production center. We further discovered that the COPD patients developed atypically in Broca’s area since they had reduced myelination and impaired microstructural integrity as measured by T_1_. The data showed that more severe dyspnea was associated with less well-developed microstructure in Broca’s area and reduced activation.

We observed no significant differences in the baseline-evoked brain activities or cognitive assessment between the two participant groups, indicating that our findings were closely related to speech production. This might be possibly due to the mild-to-moderate severity of lung disease in our patients.

Altogether, our study has demonstrated that lungs may function to shape Broca’s area as the speaking center, and this is consistent with the recent findings from the animal model research into the lung–brain axis^12,16^.

## Methods

### Participants

Thirty-five native Chinese participants (35 males, ages from 53 to 65 years) participated in this study, including 18 chronic obstructive pulmonary diseases (COPDs) and 17 healthy controls (HCs), with normal or corrected-to-normal vision. Each participant completed a series of assessments, including a basic demographic questionnaire, the Modified Medical Research Council (mMRC) Dyspnea Scale to evaluate the severity of breathlessness, and the Montreal Cognitive Assessment (MoCA) for cognitive function evaluation. To ensure the validity of the study, all participants were carefully screened to exclude those with a history of substance abuse, other pulmonary diseases, or neurological disorders, which could confound the results. Participants with COPD were included if they had a clinical diagnosis of mild to severe COPD for over one year, based on symptoms and previous pulmonary function tests, and had no other pulmonary diseases. The diagnostic criteria followed the Global Initiative for Chronic Obstructive Lung Disease (GOLD) guidelines: a post-bronchodilator ratio of forced expiratory volume in one second (FEV1) to forced vital capacity (FVC) of < 70%, and FEV1 ≥ 30% of the predicted value. This study was approved by the ethics committee at the Shenzhen Institute of Neuroscience, and all the participants provided informed consent for their participation. Detailed demographic information was listed in Table S1.

### Stimuli and Experimental Design

The study included two experimental tasks: the Noun Chinese Character (e.g., “花” meaning “flower”) and the Verb Chinese Character tasks (e.g., “踢” meaning “kick”). In both tasks, participants were instructed to overtly name the presented Chinese characters as quickly and accurately as possible. In the baseline condition, participants were required to maintain focus on the presented non-character symbols without verbal response (see Figure S2). All Chinese characters were selected from first-grade Chinese language textbook to ensure familiarity among participants. The experiment used a blocked design with five blocks each for the Noun and Verb tasks and nine blocks for the baseline condition. Each experimental block contained eight trials, and each baseline block contained nine trials. During each trial, a stimulus (e.g., character or symbol) was displayed for 0.5 seconds, followed by a 2.5-second blank interval. The stimuli within each block were presented in a pseudo-randomized order to minimize predictability and reduce potential biases. This design aimed to ensure a balanced and controlled presentation of stimuli across conditions.

### MRI Data Acquisition and Preprocessing

T1-weight structural and fMRI data were collected on a Siemens 3.0 Tesla MRI scanner using 64-channel head coil. High-resolution anatomical images were acquired using a T1-weighted, three-dimensional gradient-echo sequence (TR = 2300ms, TE = 2.07ms, Flip angle = 7°, Voxels = 0.8mm × 0.8mm × 0.8mm). The fMRI data were acquired through Echo Planar Imaging Sequences (TR = 2000ms, TE = 30ms, Flip angle = 90°, Voxel size = 3.5mm × 3.5mm × 3.5mm, Volumes = 211, FOV = 224mm, acquisition matrix = 64 × 64, slice thickness = 3.5mm). The visual stimuli were presented using a liquid crystal display projector and back-projected onto a screen positioned at the rear of the scanner bore. Participants viewed this projection via a mirror attached to the 64-channel head coil. Prior to each functional scan, 6 seconds of dummy scans were conducted to facilitate T1 equilibration.

The preprocessing of all images was carried out using the SPM12 (http://www.fil.ion.ucl.ac.uk/spm/software/spm12) and DPARSF (http://rfmri.org/DPARSF) toolboxes. The T1-weighted images were segmented into white matter, gray matter, and cerebrospinal fluid using DARTEL implemented in DPARSF to avoid signal confusion among these three tissue types^37-39^. For fMRI data, preprocessing steps included removing the first 3 images, slice timing correction, realignment, co-registering T1 images to the functional images, normalizing to MNI space using DARTEL, resampling to 3mm × 3mm × 3mm voxels, and applying 6-mm FWHM Gaussian smoothing. Particularly, participants with a maximum head motion greater than 3mm or a maximum rotation greater than 3° were excluded from this study. Finally, thirty-two participants (32 males, 53 - 65 years) were selected (detailed information see Table S1) for further fMRI analyses.

### Quantitative MRI Data Acquisition and Preprocessing

The parameters of quantitative MRI (qMRI) were determined according to the protocol of previous studies^33,40^. The MRI data were acquired using a 3T Siemens Magnetom Prisma scanner (Siemens Healthcare, Erlangen, Germany) equipped with a 64-channel head coil. Quantitative measurements of MTV and T1 were obtained from spoiled gradient echo (SPGE) images with flip angles of 4°, 10°, 20°, and 30° (TR = 12ms, TE = 2.41ms), at a voxel size of 1 × 1 × 1 mm^3^. Additionally, five spin-echo inversion recovery (SEIR) images were scanned using an echo-planar imaging (EPI) readout, a slab inversion pulse, and spectral fat suppression to correct for field inhomogeneities. The images were captured at inversion times of 50, 200, 400, 1200, and 2400ms (TR = 3000ms, TE = 49ms), with a voxel size of 2 × 2 × 7 mm^3^. Among them, two participants were registered and resampled with other participants due to the inconsistent slice thickness. Finally, 30 participants were adopted for follow-up qMRI analysis (detailed information see Table S1).

## Data Analysis

For individual-level analysis, All individual participant t-maps (i.e. Noun Chinese Character vs. Baseline, Verb Chinese Characters vs. Baseline, Noun Chinese Character vs. Verb Chinese Characters, Verb Chinese Characters vs. Noun Chinese Character, and Noun Chinese Character + Verb Chinese Characters vs. Verb Chinese Character) were generated using the general linear model based on the smoothed data, in which time series were convolved with the canonical hemodynamic response function and were high-pass-filtered at 128s. Head motion parameters of six dimensions from realignment were modeled as regressors of no interest^41^.

The contrast images subsequently analyzed at the group level using a second-level model. Specifically, the one-sample t-test was performed to generate whole-brain activation maps by averaging across all participants in each group between experimental (Noun Chinese Character and/or Verb Chinese Character) vs. baseline condition. To assess the impact of COPD on brain activation, two-sample t-test was further adopted to contrast brain activity between two groups. The significance level threshold was set at *P* < 0.05 (False Discovery Rate correction, FDR)^41^ with a spatial extent larger than 10 voxels^35^. In addition, we calculated the t-map for each participant under the baseline condition and then compared the brain activities between the two groups, applying a significant threshold of *P* < 0.05 (FDR-corrected) and *P* < 0.001 (uncorrected). We found no reliable differences across the two groups. This indicated that our observed brain activations were attributable to the speaking task.

Based on the contrast image of the two groups of participants (HC > COPE) in contrast condition (experimental versus baseline conditions), regions of interest (ROIs) were extracted by intersecting the activation map (FDR-corrected, *P* < 0.05, spatial extent >10 voxels) with the Brodmann map. To ensure comparability, the Brodman mask from DPABI was resampled to match the spatial resolution of the activation maps. To quantify the effect size of the observed differences in brain activation between the COPD and HC groups within the identified ROIs, Cohen’s d was calculated for each ROI based on the T-values derived from the second-level group analysis. The formula for Cohen’s d based on T-value is as follows:

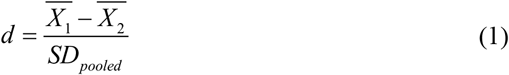

Where 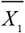 and 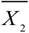 represents the mean activation values for the COPD and HC groups, respectively; *SD*_*pooled*_ is the pooled standard deviation across the two groups. The formula for independent samples t-test is as follows:

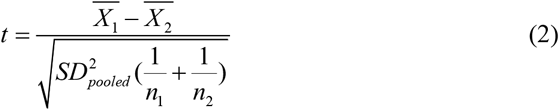

where n_1_ and n_2_ represents the sample sizes for the COPD and HC groups, respectively. Combing the formula of (1) and (2), the final formula to calculate Cohen’s d from the t-value is:

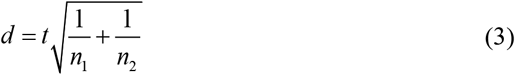

To assess the differences in brain region activation between the COPD and HC groups, we selected a series of literature-based regions which were reported to related to the lung function, including the bilateral pars opecularis (MNI: -54, 8, 16; 50, 10, 18)^28^, the left inferior frontal triangularis (MNI: -50, 36, 22)^29^, the left inferior frontal gyrus (MNI: -30, 50, 20)^30^, the left middle frontal gyrus (MNI: -40, 40, 16)^29^, the right pre-supplementary motor area (MNI: 4, 12, 56)^28^, the left inferior precentral gyrus (MNI: -52, -8, 50)^31^, the left middle insula (MNI: -42, 2, 2)^30^ and the right insula (MNI: 38, 10, 8; 45, 16, -2)^32^. After extracting activation values at specified MNI coordinates from these regions, we performed a two-sample t-test to compare the activation between the two groups. The significance level threshold was set at *P* < 0.05 (Benjamini-Hochberg FDR correction).

The SPGE and SEIR images were processed using the mrQ toolbox (https://github.com/mezera/mrQ) in MATLAB to generate participant-specific T_1_ maps^35,40^. T_1_ maps provide voxel-wise longitudinal relaxation times, adding in the characterization of tissue properties like myelination and fluid content. To focus on Broca’s area (left BA44), a mask was extracted from the Brodmann atlas and aligned with each participant’s T_1_ map. The mask was resampled to ensure spatial consistency across participants, and mean T_1_ values for Broca’s area were calculated. A non-parametric rank sum test was employed to compare T_1_ values between two groups, as data distribution did not meet normality assumptions. Additionally, the Cohen’s d values (1) of mean T_1_ values were further evaluated to quantify the effect size of between two groups within the Broca’s area. We further computed the partial correlations between mean T_1_ values and mean values of the top 30% activation values in the left BA 44, controlling for the education level.

Finally, we calculated Pearson’s correlation coefficient to evaluate the relationship between modified Medical Research Council (mMRC) Dyspnea Questionnaire scores and activation values in the identified regions of interest (ROIs), as well as the mean T_1_ values in the left BA 44, where the correlation was deemed significant at a threshold of *P* < 0.05. Higher mMRC scores indicate more severe breathing symptoms.

## RESOURCE AVAILABILITY

### Lead contact

Further information and requests should be directed to and will be fulfilled by the lead contact, Li-Hai Tan.

## Data availability

Data for which patients have consented to public release will be made available at a specified website by the journal, e.g., the Data Archive for the BRAIN Initiative (DABI; https://dabi.loni.usc.edu).

## Code availability

The analysis and data visualization code will be made available in the Shenzhen Institute of Neuroscience website upon publication.

## ACKNOWLEDGEMENTS

We thank Yujie Liu, Ruijie Zhou, Yulong Zhou, Wenqi Weng, Jinjian Wu, Luxia Yang, and the physicians of the Respiratory Department of First Affiliated Hospital of Guangzhou University of Chinese Medicine for their assistance in recruiting the patients. We also thank Min Xu for constructive comments. This study was supported by the STI 2030—Major Projects (grant no. 2021ZD0200500).

## AUTHOR CONTRIBUTIONS

S.Y.C., S.J.Q. and L.H.T. conceived the study. S.Y.C., W.W.Z., Y.Q.H. S.J.Q. and L.H.T. designed the experiment. S.Y.C., W.W.Z., and Y.Q.H. collected the data and performed the data analyses. S.Y.C., W.W.Z., Y.Q.H. and L.H.T. prepared the manuscript. All authors reviewed and edited the manuscript. L.H.T. provided supervision and funding for all aspects of the study.

## DECLARATION OF INTERESTS

The authors declare no competing interests.

## Additional information

The work contains Supplementary information.

## Supplementary information

**Supplementary TABE 1.**
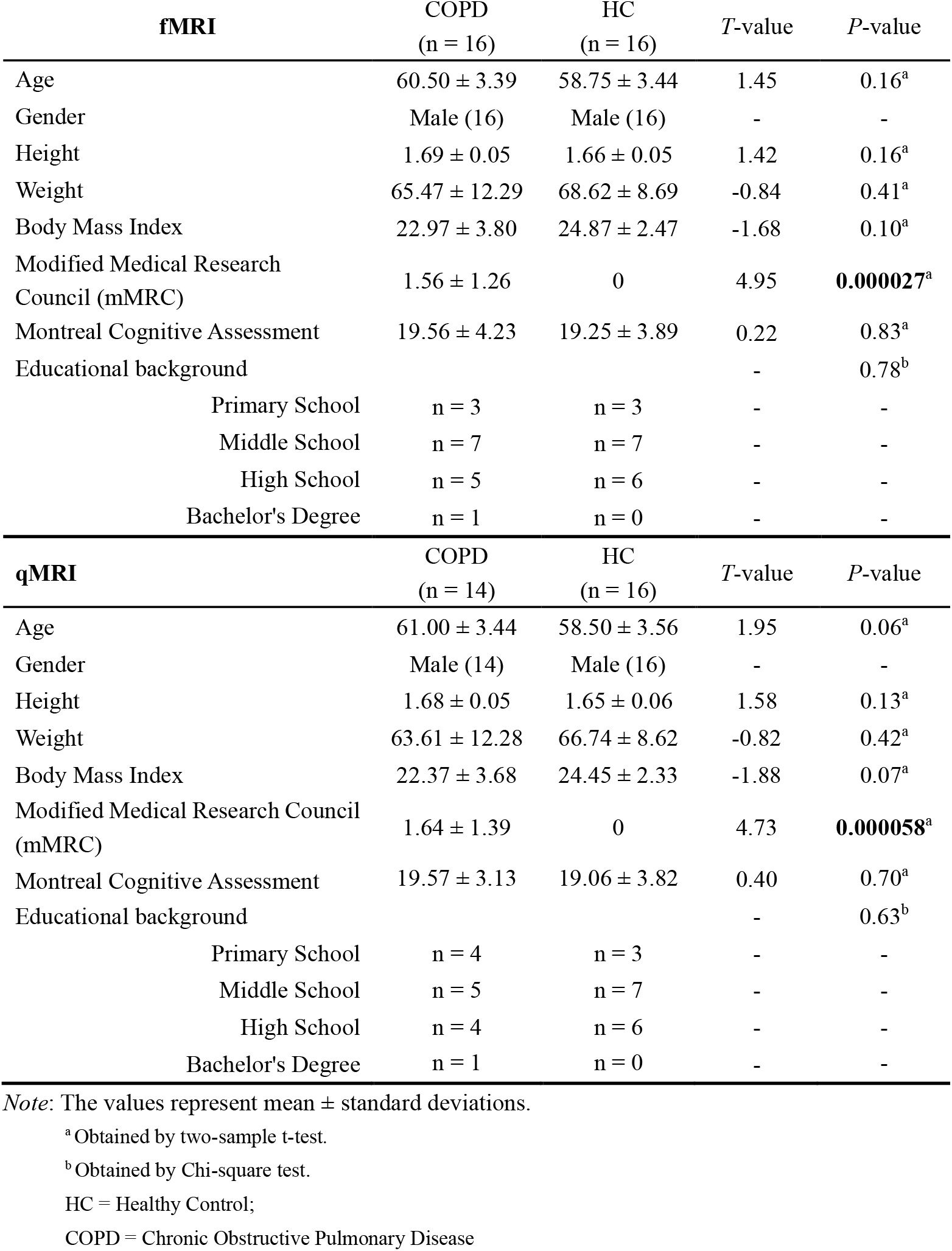
Demographic information of the participants.

**Supplementary TABLE 2.**
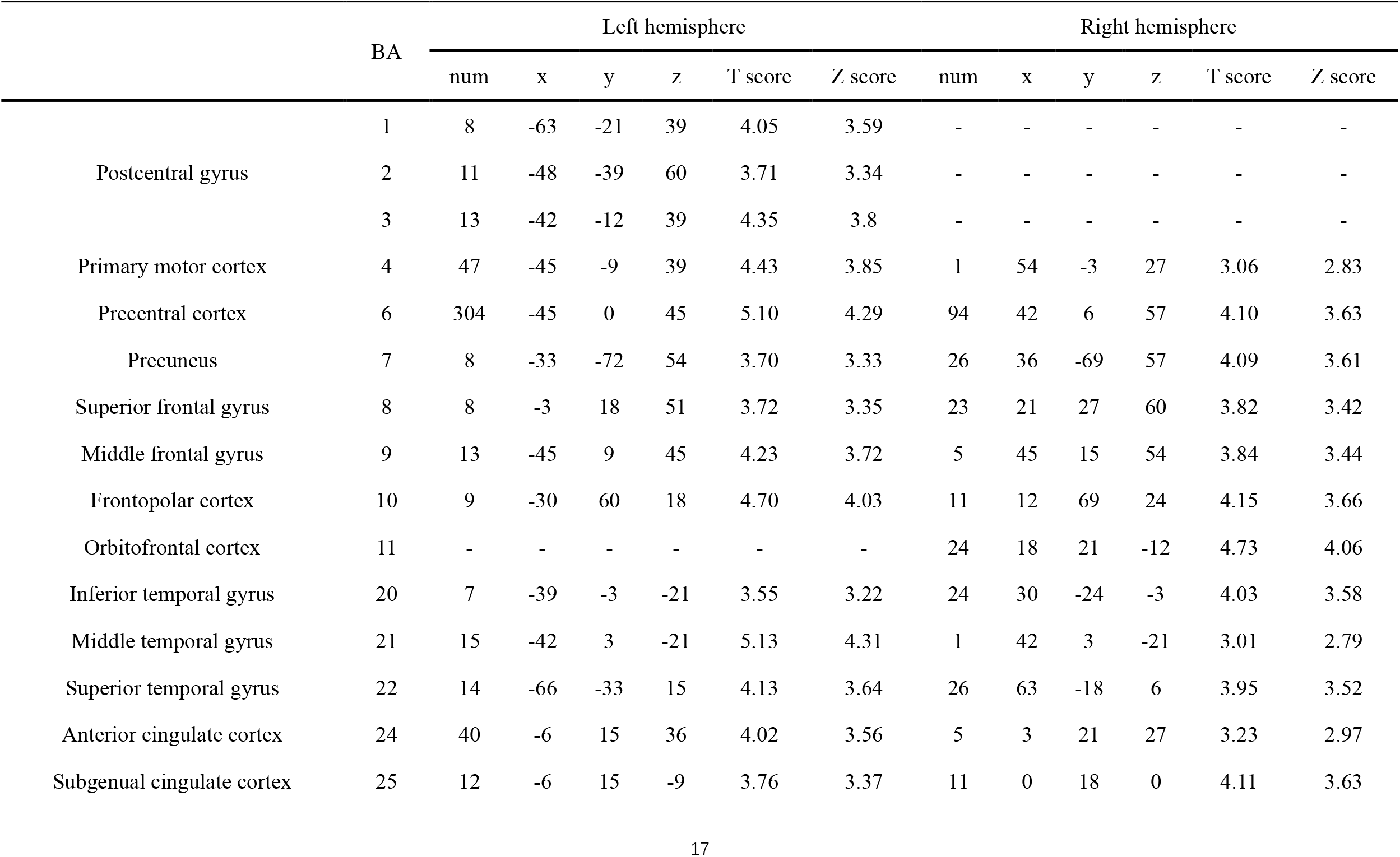
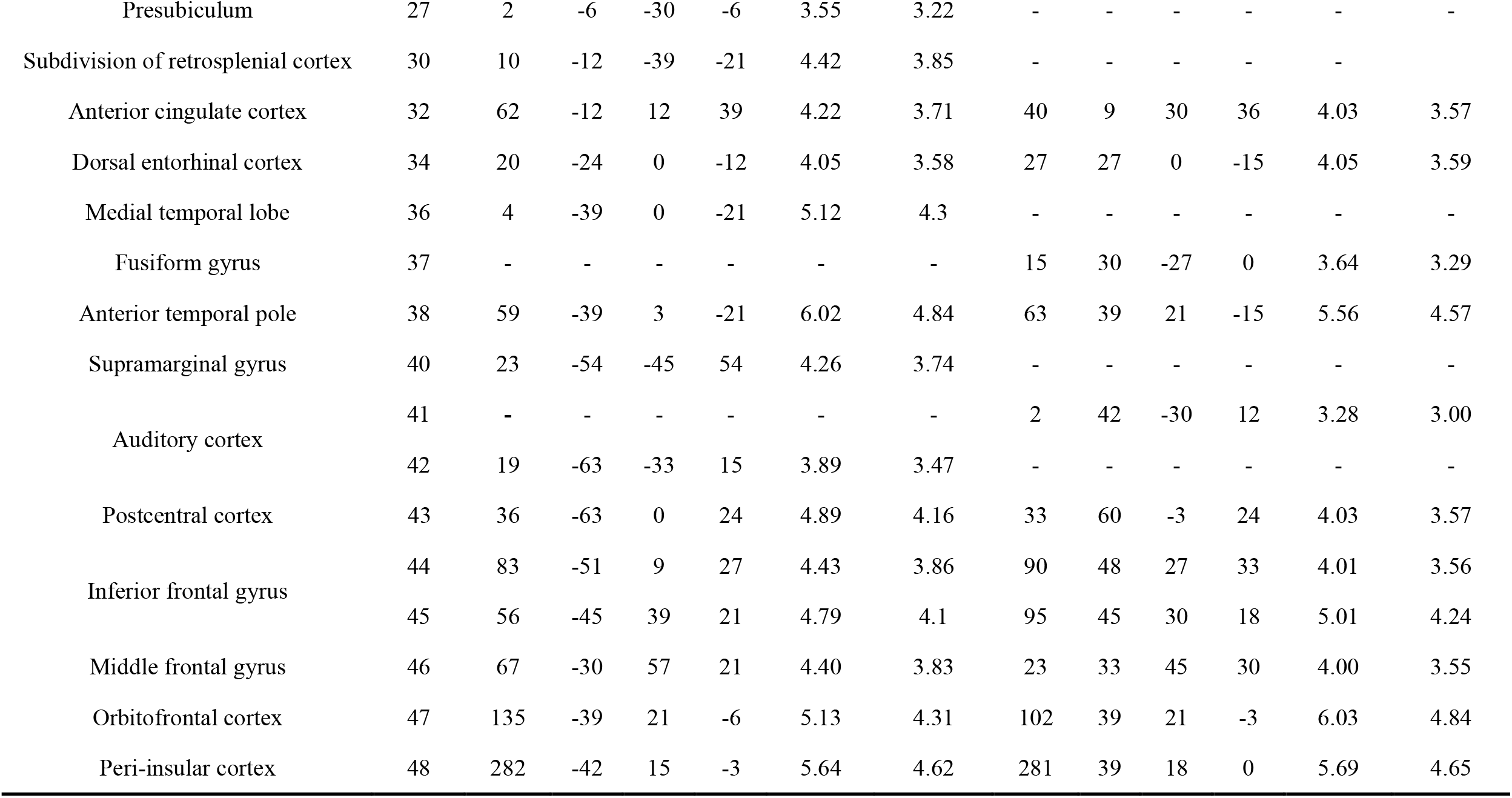
MNI Coordinates of activation peaks.

**Supplementary FIGURE 1.**
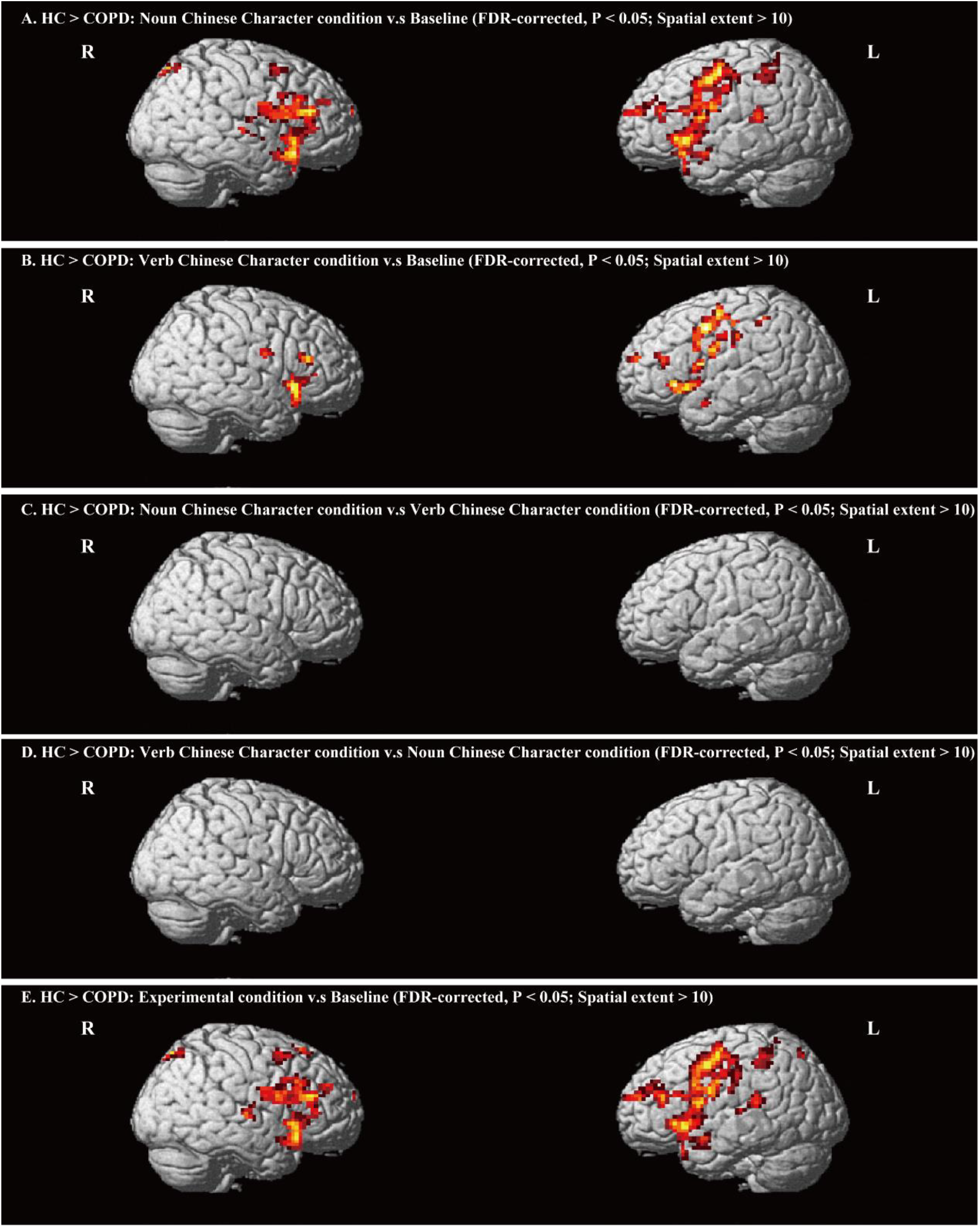
Statistical parametric maps of group activation differences between tasks (FDR-corrected *P* <0.05 with spatial extent n≥10 voxels).

**Supplementary FIGURE 2.**
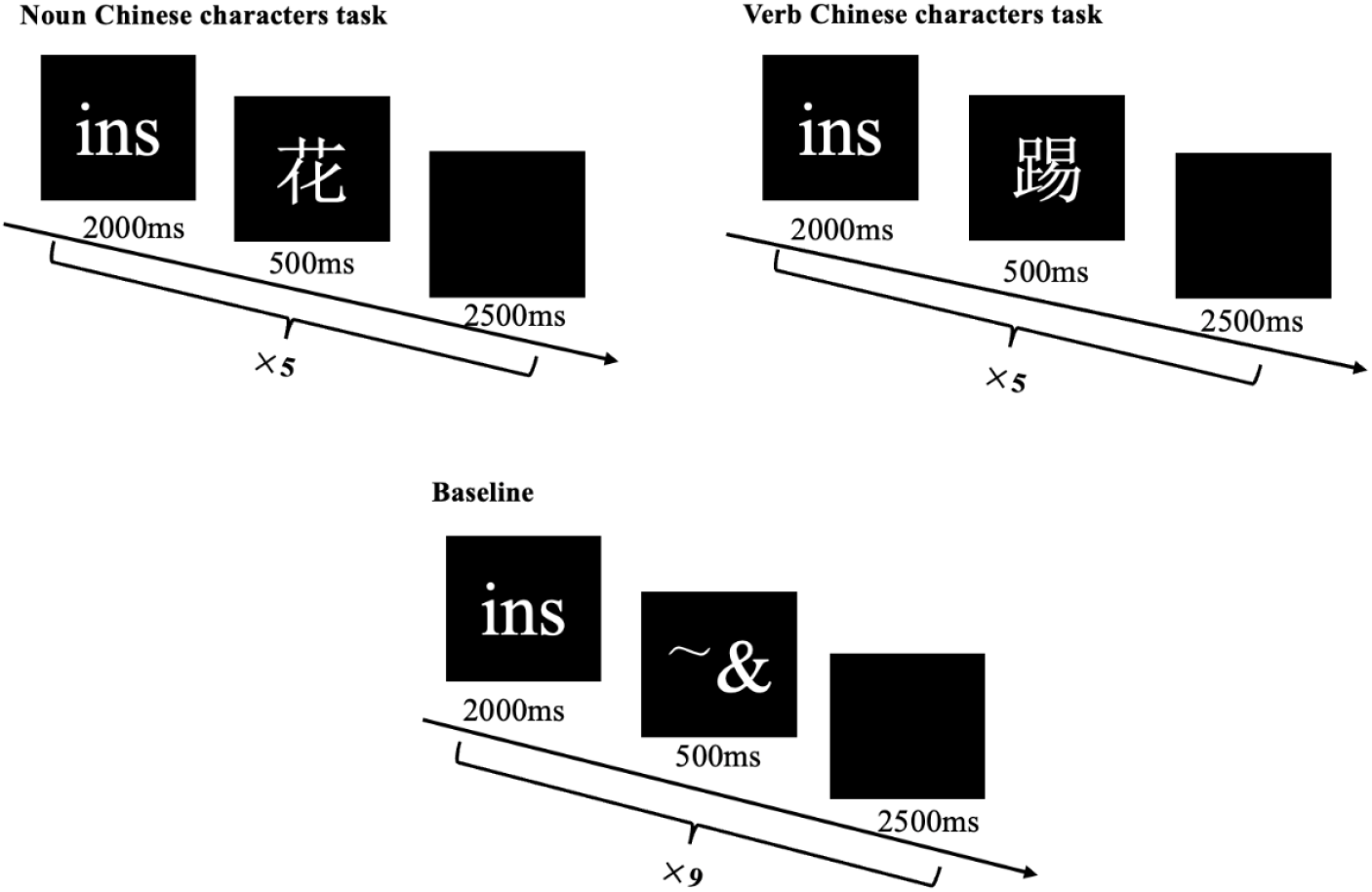
fMRI Paradigm and Stimuli.

## Notes

### Competing Interest Statement

The authors have declared no competing interest.

### Summary of Updates

Modified some language expressions and added the correlation results between the activation value and T1 value under baseline conditions ('By contrast, in the baseline condition, no significant partial correlation was observed between the top 30% activation values and mean T1 values (r = -0.0052, P = 0.9799), suggesting that the observed negative correlation in the experimental condition is specific to task-related activations.') revised the order of author first name and last name

